# Derivation of Steady-State First-order Rate Constant Equations for Enzyme-Substrate Complex Dissociation, as well as Zero-order Rate Constant Equations in Relation to Background Assumptions

**DOI:** 10.1101/2022.12.15.520621

**Authors:** Ikechukwu I. Udema

## Abstract

The maximum velocity (*V*_max_) of catalysis and the substrate concentration ([*S*_T_]) at half the *V*_max_, the *K*_M_, are regarded as steady-state (SS) parameters even though they are the outcomes of zero-order kinetics (ZOK). The research was aimed at disputing such a claim with the following objectives: To: 1) carry out an overview of issues pertaining to the validity of assumptions; 2) derive the needed steady-state (SS) equations distinct from Michaelian equations that can be fitted to both experimental variables and kinetic parameters; 3) calculate the SS first-order rate constant for the dissociation of enzyme-substrate complex (ES) to free substrate, S and enzyme, E; 4) derive the equation of rate constant as a function of the reciprocal of the duration of each catalytic event in the reaction pathway. The experimental values of the data were generated by Bernfeld and Lineweaver-Burk methods. The calculated SS 1^st^ order-order rate constant was « the zero-order Michaelian value, and the difference is ≈ 97.59 % of the zero-order value; the SS catalytic rate differed from the zero-order catalytic rate by ≈ 76.41 % of the latter value; and it was ≈ 93.87 % with respect to the 2^nd^ order rate constant for the formation of enzyme-substrate complex. The equations of time-dependent rate constants, *K*_M_, and dissociation constants were derived. The concentration [*S*_T_] of the S must be > the concentration ([*E*_0_]) of the E for the quasi-steady-state assumption (or approximation) to hold. The SS kinetic parameters are not equivalent to zero-order parameters.

## 1. INTRODUCTION

It is well known that events within the catalytic site during the first enzymatic turnover are best studied during the steady-state burst phase [1]. Such events cannot preclude different rate constants and other Michaelian parameters in the light of what are regarded as steady-state parameters [1]. With reference to the literature [1], Sassa *et al*. [2] noted three standard kinetic approaches for characterising the behaviour of a chosen enzyme model. Such approaches are: 1) steady-state; 2) pre-steady-state; and 3) single-turnover. Before proceeding further, the definition of turnover is: The turnover number represents the maximum number of substrate molecules converted to product per active site per unit time, or the number of times the enzyme is “turned over” per unit time [3]. However, it is imperative for the researcher to establish appropriate underlying assumptions as long as any of the approaches need to be adopted. Thus, whenever the Michaelis–Menten equation is to be used to estimate Michaelian parameters, the Michaelis– Menten constant, *K*_M_, and the maximum velocity of product formation (catalytic rate), *V*_*max*_, it is essential to know whether or not the steady-state assumption is valid in any given experimental assay for an enzyme catalysed reaction [3]. There seems to be more emphasis on catalytic efficiency and proficiency in kinetic investigation of enzymatic action. Thus, the emphasis is on interpreting the steady-state kinetic parameters in terms of enzyme structure and individual steps in the reaction pathway. Hence, according to Johnson [1] (with reference to the literature [4]), there is justification in the choice of *k*_cat_/*K*_M_ rather than *K*_M_ and *k*_cat_ as a primary kinetic parameter. *k*_cat_/*K*_M_ is regarded as the most important as it quantifies enzyme specificity, efficiency, and proficiency [4]. The parameters are regarded as steady-state parameters [1].

One of the goals of this research is to go beyond the standard rate constants, the second-order rate constant for ES formation and the catalytic rate, *k*_cat_, for the formation and release of product, P. This is notwithstanding the fact that the rate constant for the dissociation of the enzyme-substrate complex (ES) to free substrate (S) and free enzyme (E) is a regular feature in the Michaelian constant. Another goal is to examine the ascription of Michaelian parameters— the kinetic parameters in general—to steady-state characteristic parameters. This means that the pre-steady-state (or burst) phase, steady-state, and zero-order state (or zone) all have their own rate constants. None of the phases or stages (zones) is permanent; the long-lasting zone (the zero-order zone) soon disappears when substrate depletion takes centre stage, and in particular if there is a way of evacuating the product without product inhibition at any time, as in an *in vivo* or system scenario. The research is intended to show qualitatively (i.e., without any calculation for now) that the apparent rates are the sum of the reciprocal of each of the durations of various stages of the reaction pathway.

The primary goal of this study is to determine the steady-state first-order rate constant for the dissociation of ES to E and S in relation to the background assumptions, which must also be valid in order to validate the enabling kinetic parameters. Recall that *K*_M_ can be described as a mixed “pseudo-equilibrium constant” in that it is = *k*_-1_/*k*_1_ + *k*_cat_ /*k*_1_ (*k*_-1_, *k*_cat_, and *k*_1_ are the 1^st^ order rate constants for ES → E + S, the 1^st^ order rate constant for product formation release, and the 2^nd^ order rate constant for enzyme-substrate ES formation, E + S → ES). However, a preprint report [5] has called the Michaelis-Menten constant a steady-state constant but fell short of indicating how the steady-state variables can be measured or calculated to enable the evaluation of [*E*_F_]_SS_ [*S*_T_]_SS_/[*ES*]_SS_, where SS and [*E*_F_]_SS_ stand for steady-state and the steady-state free enzyme concentration, respectively. To be conscious of the very high standard of presentation demanded by publishers also requires that questions be asked consistently. For instance, is the *K*_M_ an outcome of a linear relationship between initial rates and substrate concentrations less than *K*_M_? It is well understood that *in vivo* scenarios present a dynamic scene in which products are taken away for the next stage in the metabolic pathway, causing the “equilibrium” to shift too far to the right, such that what dominates in an earlier step or stage is the *k*_cat_ with very low (or nonexistent) *k*_-1_. In other words, for such reactions, *k*_−1_ is negligible in front of *k*_cat_, and in consequence, for a given *k*_cat_, the higher the *k*_1_, the lower the *K*_M_ and the higher the flux provided by the catalytic step. In other words, fluxes depend on *k*_1_ in many in vivo physiological conditions [5].

In an industrial situation, optimisation of product formation occurs by exiting the products as they are formed. However, control demands a way of regulating product formation, as the case may be, by whatever principle, Le Chatelier’s principle, for instance. *In vivo*, or the true system scenario, requires that as soon as an individual responds to the need for a prandial diet, the system mobilises the needed enzymes for the digestion of proteins, for instance, yet gradually ceases as soon as a substantial amount of digestion has been achieved. Under normal circumstances, sufficient blood sugar must inhibit the enzymes needed for the release of glycogen. It is not just a function of the hormonal and nervous systems alone. Simulation of such processes *in vitro* may be speculative for now. As long as determining *k*_-1_ is of primary interest, the following objectives are pursued: 1) conduct an overview of issues pertaining to the necessary condition of the validity of assumptions; 2) derive required steady-state (SS) equations distinct from Michaelian equations that can be fitted to both experimental variables and kinetic parameters; 3) calculate the first-order rate constant for the process, ES → E + S, peculiar to the steady-state zone of an enzyme catalysed reaction pathway; 4) derive the equation of rate constant as a function of the reciprocal of the duration of each catalytic event in the reaction pathway.

## 2. THEORY

In this section, all the conditions for the validity of the steady-state assumption are examined. Here, two types of assumptions are analysed and the equations to be fitted to the experimental variables are derived. Time-dependent rate constants specific to different zones of the kinetic cycle (or, more precisely, reaction pathway) are to be derived and investigated in order to deduce mutually dependent opposing first order rate constants.

### 2.1 Assumptions and background condition for validity

It has been shown that the standard quasi-steady-state assumption (approximation), sQSSA, holds if the initial substrate concentration [*S*_T_] is much larger than the total or initial enzyme concentration [*E*_T_] [6]. They refer to a “brief transient,” *t*_c_ (the time of ES buildup), which is indeed very short compared to *t*_s_ (the time during which the substrate concentration changes appreciably), *i*.*e*.*t*_*c*_ « *t*_s_. The equations for the two time scales are [8]:

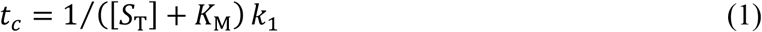

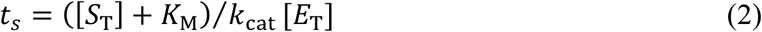

In terms of time regime, the sQSSA may hold if *t*_*c*_ « *t*_s_, but it needs to be stated that this may also be contingent on the nature of the substrate, which can influence the magnitude of the Michaelis-Menten constant, *K*_M_ regardless of if [*S*_T_] » [*E*_T_]. A gelatinised insoluble starch is expected to present effect different from an ungelatinized starch. A 2^nd^ criterion is such that:

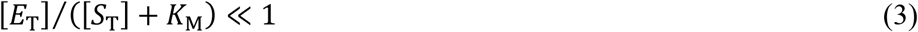

Alternatively, Eq. (3), which can be found in the work of Schnell [3], must also bear the following relationship:

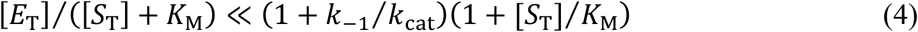

Equation (4) is attributed to Segel and Slemrod [6]

The reactant stationary assumption (RSA) proposes that during the initial transient, the concentration of substrate [*S*_(t)_] in a very short time is ≈ [*S*_T_], the initial concentration of substrate in time = 0 [3]. This cannot be different from the condition that validated the pseudo-steady-state assumption (also known as sQSSA), given that saturation of the enzyme within a defined time frame, specifically a short duration, occurs primarily at much higher [*S*_T_]. Therefore, it is not certain how the hyperbolic relation of the velocity versus [*S*_T_] can be observed if, as one of the conditions for the validity of RSA, the relationship [*E*_0_] ≈ [*S*_T_] ([*E*_0_]/[*S*_T_]) [3] is involved. Someone must realise that mass-mass concentration as opposed to mole-mole concentration may be deceptive in that the mass concentration of the enzyme, for instance, may be « the substrate concentration; but being that the molar mass of the enzyme is « the molar mass of the substrate, its molar concentration could be > the molar concentration of the substrate.

Of course, an appropriate choice of the assay duration is very relevant in order to avoid saturation even at the lower end of the substrate concentration range. This is where rapid methods, stopped flow and chemical-quench-flow are useful: In this regard, it has been observed that steady-state and transient-state kinetic studies complement one another, and steady-state analysis should be performed prior to properly designing and interpreting experiments using transient state kinetics (TSK) and its methods [1]. TSK could be defined as an initial transient kinetics because it has a very short time scale and [*S*_(t)_] ≈ [*S*_T_].

The criterion that [*E*_T_] ≈ [S_T_] for RSA to be valid requires that [*E*_T_] « *K*_M_ if the relationship [3] below holds:

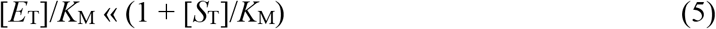

The implication of Eq. (5) is the admissible fact that steady-state kinetics demands [S_T_] » [*E*_T_] because some, if not all, values of [*S*_T_] within the range may be > *K*_M_. This explains in part why Briggs and Haldane’s [8] approach in the derivation of the Michaelian-Menten [9] equation demands that “ES need not be in equilibrium with E and S, but within a very short time after starting the reaction, the rate of formation of ES will almost balance its rate of destruction.” Hence, ES builds up to a pseudo-steady-state level where its concentration is nearly constant. [3]. This is due to the fact that as soon as the enzyme is released, it quickly binds to abundant substrate, ensuring nearly constant availability of ES, which is referred to as “one in pseudo-steady-state” to imply the following relationship:

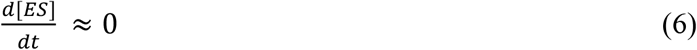

While it is proposed that [*S*_t_] need not be ≈ [*S*_T_] in order to have a valid steady-state assumption, it is clear that Eq. (6) may hold because it is the availability of S in excess of E that ensures its occurrence. Given that [ES] = *v*/*k*_cat_ and *v* = − d[*S*_T_]/d*t* /*M*_s_ (where *M*_s_ is the molar mass of the substrate), Eq. (6) can be rewritten as d*v*/d*t*/*k*_cat_, followed by:

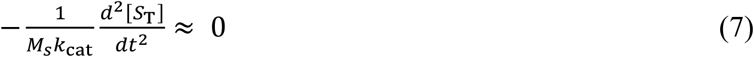

Therefore, Eq. (6) and Eq. (7) are equivalents, such that while Eq. (6) defines the pseudo-steady-state requirement, it is highly complemented by Eq. (7), which also defines the RSA requirement, in that, according to Schnell [3],

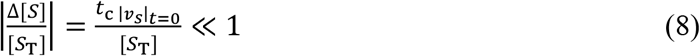

where, |*v*_*s*_|_*t*=0_, is the magnitude of the rate (velocity) of utilisation of the substrate, given as:

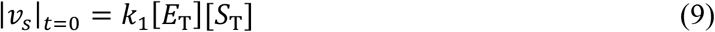

Therefore, if Eq. (7) holds, then 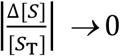 is also possible. Therefore, the view that [*S*_(t)_] ≈ [*S*_T_] need not hold [10] for steady-state assumption to be valid appears to be inconsistent with the Briggs and Haldane method, which takes into account the steady-state assumption requirement even if it is seen to be applicable to RSA because the substitution of Eq. (9) into Eq. (8) followed by the substitution of Eq. (1) leads to the same equation as Eq. (3) that defines the condition for the validity of a quasi-steady-state assumption. If [*S*_(t)_] ≈ [*S*_T_], is a valid condition that validates RSA, the same is very applicable to the steady-state assumption. What is in a quasi-steady state is the ES, which is made possible at the initial transient in short duration due to possibilities such as Δ[*S*_T_] → 0, implying that [*S*_(t)_] ≈ [*S*_T_], that is, stable concentration of S, is applicable to both QSSA and RSA.

### 2.2 Derivation of equations to be fitted to experimental variables

The equations to be derived are those for the calculation of rate constants that cannot be directly calculated based on experimental and measured data. However, contrary views in the literature [3] suggest that overestimation of Michaelian parameters may be due to outliers or error-laden experimental variables, leading to the development of the direct linear method or its reciprocal equivalent and, more recently, nonlinear regression analysis [11]. The rate constants are the lower steady-state first-order rate constant for the dissociation of ES into free E and S, the lower catalytic rate constant, the lower second-order rate constant for the formation of ES, and consequently, the steady-state dissociation constant.

#### 2.2.1 Derivation of first-order rate constant for the decomposition of ES into E and S

It has been shown in a previous publication [12] that the concentration of free enzyme can be given as:

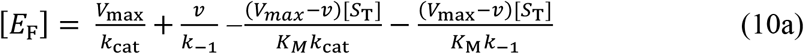

where, [*E*_F_], [*S*_T_], *k*_cat_, *V*_max_, *v, K*_M_, and *k*_−1_ are the concentrations of the free enzyme, and substrate, catalytic 1^st^ order rate constant, maximum velocity of enzymatic action, velocity of catalysis, Michaelis-Menten constant, and the post steady-state 1st order rate constant for the process, ES → E + S. Taking the issue of what is regarded as steady-state kinetic constants to the forefront, there should be a critical need to appraise Eq. (10a), whose validity has been attested to in the literature [12]. In the first place, *v* can take any value between 0 and *V*_max_/2. If so, one may wish to know if *V*_max_ and its associated *K*_M_ are measured at the steady-state zone; if so, is the zero-order zone in which rate is independent of substrate concentration the same as the steady-state zone where a linear relation exists between *v* and [*S*_T_]? The Michaelian principle requires that the substrate concentration be sufficiently > enzyme concentration such that, within the duration of the assay, all the molecules of enzyme will always be complexed with the substrate but not ad infinitum; this implies that, with the abundance of the substrate, as soon as the product is released, the free enzyme is closely exposed to the substrate to which it binds. On account of the foregoing, Eq. (10a) is restated as:

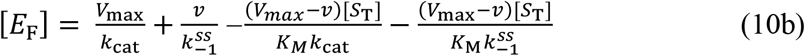

where 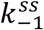 is the steady-state 1^st^ order rate constant for the dissociation of ES into E and S.

Within a defined life span of the zero-order zones, an increase in substrate concentration will not increase the rate of catalysis. Again, is the enzyme saturated with the substrate at the steady-state zone? It should be noted that saturation is a time-dependent phenomenon, so any transient time scale (« milliseconds) duration of the assay does not provide enough time for all of the enzymes to become saturated, even if [*S*_0_] » [*E*_0_] and stirring of the reaction mixture are active. Meanwhile, if *v* = *V*_max_/2 in Eq. (10a), then the concentration of the substrate should be equal to the *K*_M_. Consequently, Eq. (10a) simplifies to [*E*_F_] = *V*_max_/2*k*_cat_, which means that the concentration of the free enzyme is equal to one-half of the total enzyme concentration, [*E*_T_]. Thus, the *V*_max_ is not achieved outside the enzyme’s saturation zone, though it may be calculated if the *K*_M_ is known, giving a value similar to the experimental value. The steady-state zone does not yield the main kinetic parameters (Michaelian parameters) such as *K*_M_ and *V*_max_.

Once again, bearing in mind that a steady-state scenario is of interest, the following equation should hold:

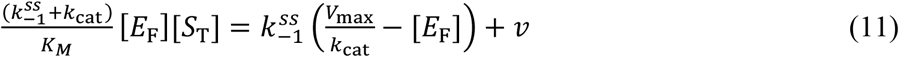

(*V*_*max*_/ *k*_*cat*_) − [*E*_F_] = [*ES*] Therefore, Eq. (1) can be written as:

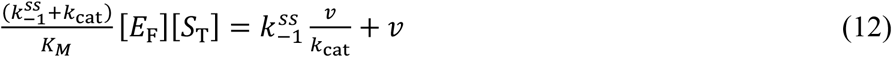

Substituting Eq. (12) into Eq. (10) gives:

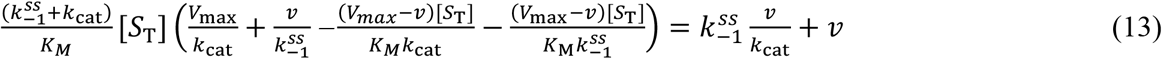

Rearranging the right hand side and eliminating 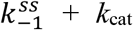 gives after rearrangement the following.

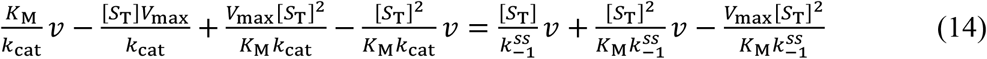

At this point, the rate of product (P) formation must be changed to the rate of substrate disappearance (S), keeping in mind that d[P]/dt is not considered to be **≈** 0. Given that *v* = − d[*S*_T_]/d*t*, Eq. (14) can be rewritten as follows:

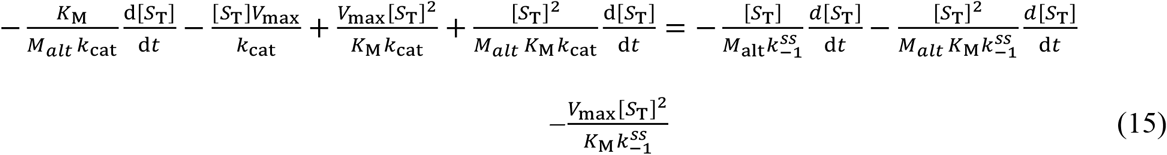

Here, the molar mass (*M*_alt_) of maltose is introduced to account for the fact that the change in mass of substrate due to enzymatic conversion to product is, in line with mass conservation law, approximately equal to the yielded mass of maltose being converted to a molar unit. Dividing both sides of Eq. (15) by [*S*_T_] followed by cross multiplication by d*t* gives:

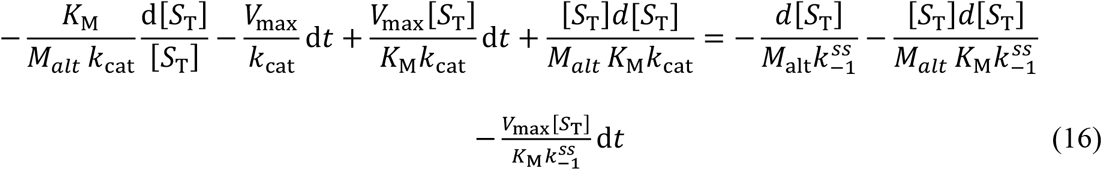

Integration of Eq. (16) gives:

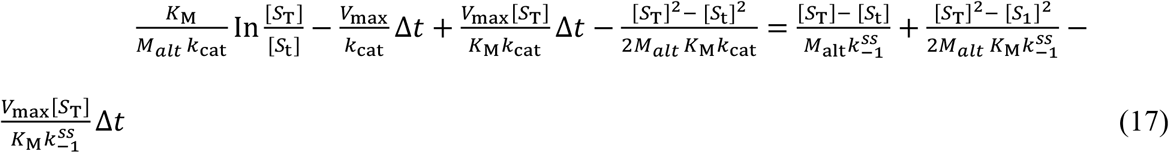

where, Δ*t* and [*S*_t_] are the duration of assay and substrate concentration at the end of the duration of assay. Besides, [*S*_t_] ≅ [*S*_T_] − *M*_alt_ *v* Δ*t*; and dividing through by Δ*t* and rearrangement gives:

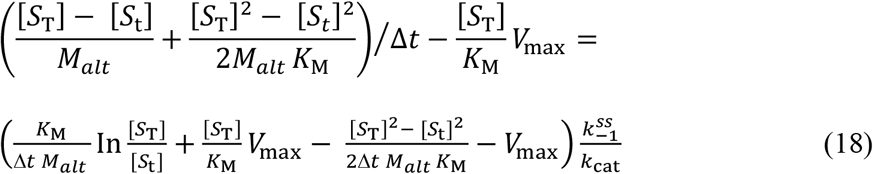

A plot of the left hand side versus the right hand side gives the slope, 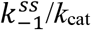.

Let us recall that the quasi-steady-state assumption for the ES expressed as Eq. (6) is the outcome of Briggs and Haldane’s [8] exposition. This calls for an experimental design that permits the measurement of reaction velocities (the so-called initial rates) in regimes where the rate of formation of the product, P, is directly proportional to [*S*_0_], that is, the rate of formation of P is linear with time. Yet *k*_cat_ (or *V*_max_) is not measured at the zone of the reaction pathway wherein the d[P]/dt is proportional to [*S*_0_]; it is a function of zero-order kinetics, which is substrate concentration invariant even if it is used to derive the Michaelis-Menten equation. Based on the assumption that the conversion of E and S to ES is faster than the decomposition of ES into E and P (*k*_cat_ « *k*_−1_), the equation [*ES*] = [*E*][*S*]/*K*_M_ is derived. The rapid equilibrium assumption was postulated by Michaelis-Menten; on the other hand, if *k*_cat_ is » *k*_−1_, the Van Slyke-Cullen [13] scenario is the case that gives rise to the Van Slyke-Cullen constant, *k*_cat_/*k*_1_. Meanwhile, the value of *V*_max_ (*k*_cat_ [*E*_0_]) can also be regarded as an asymptotic value, achieved when the zero-order rates are attained such that the initial velocity transits from a linear relationship with the substrate concentration to a “zero-order relationship” with the substrate concentration. This is totally not applicable to a steady-state zone, which is substrate concentration-dependent. As posited elsewhere [14], the measurable quantities [*E*_0_] and [*S*_T_], but not to the exclusion of the *V*_max_ and *K*_M_, which are experimentally determined, are explored for the determination of the immeasurable; this enhanced the derivation of the original Michaelis-Menten equation.

It is also important to recognise that there is skepticism about steady-state kinetics, which, while useful, is characterised by a scarcity of information on catalysis and information content on rate constants [14]. Yet the same author [14] is of the view that the design of pre-steady-state and single-turnover kinetic measurements depends on the estimates of the lower limits of *k*_cat_ and *k*_cat_/*K*_M_ from steady-state measurements. This implies that there may be upper limits to the quantitative values of the former; such cannot be found outside the mixed zero-order zone, let alone outside the zero-order zone. Similar to an earlier view, steady-state measurements are characterised by multiple turnovers under conditions that satisfy the relationship: [*E*_0_]/(*K*_M_ + [*S*_0_]) «1, such that the time courses are linear under the condition of an initial velocity far from the asymptotic value of the velocity of catalytic action. The simple deduction is that *V*_max_ (or [*E*_0_] *k*_cat_) and *K*_M_ parameters are not steady-state parameters; the “big” question is, then, how can steady-state kinetic parameters be determined?

#### 2.2.2 Derivation of steady-state second-order rate constant for the formation of ES and catalytic rate constant for the formation of product

Equation (18) enables the determination of the steady-state value of 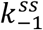 (the steady-state version of the first-order rate constant for the backward reaction of ES to give E and S in a post-steady-state zone). Also, the steady-state velocity is higher than the initial velocities, *v*, and the corresponding catalytic rate, postulated to be < upper limit value achieved at saturation, is designated as 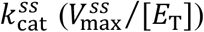. The main problem illustrated in Eq. (18) is that zero order zone (or saturation) kinetic parameters are used to calculate the steady-state first order rate constant for the backward dissociation of ES to free E and free S. The implication is that there should be a steady-state, second-order rate constant for the formation of ES, which can be determined by exploring the equation given in the literature [12]. The equation is:

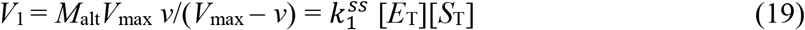

where *V*_1_ and *M*_alt_ are the velocity of ES formation and molar mass of maltose. The resulting slope from the plot of the left hand side versus [*S*_T_] gives a slope, Š_Ĺ_. The slope is further given as:

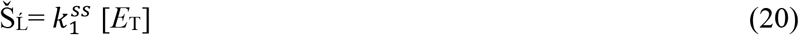

The 2^nd^ order rate constant for the formation of ES under a steady-state scenario is given as:

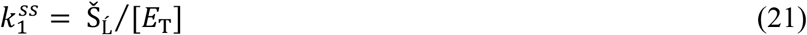

As explained in the literature, the second-order rate constant in L /mol. min for the formation of ES is related to the molar mass of the product as follows:

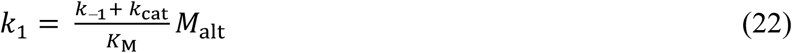

However, for the purpose of this research, Eq. (22) is restated as:

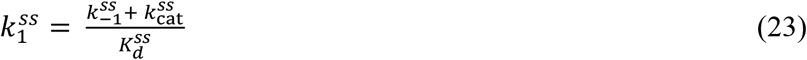

Here, 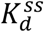 (dissociation constant) whose unit should be L/mol. min is defined as in the equation of linearity given as 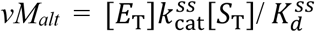. Thus, a plot of *vM*_alt_ versus [*S*_T_] gives a slope symbolised as S_§_ and defined as:

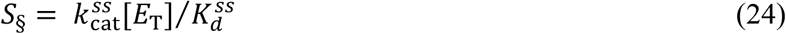

Meanwhile, Eq. (23) can be expanded to give:

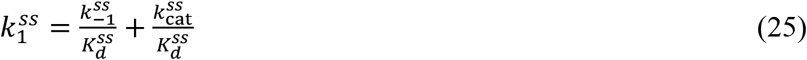

However, from Eq. (24) is given 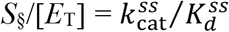. Then substitution into Eq. (25) and making 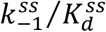 subject of the formula gives:

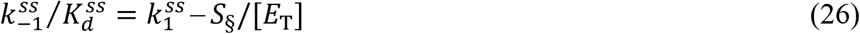

The parameter, 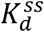 in Eq. (26) is then given as:

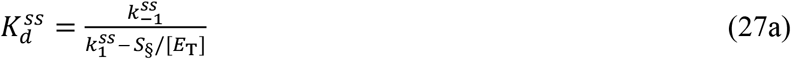

Also, the steady-state second-order rate constant for the formation of ES is given as:

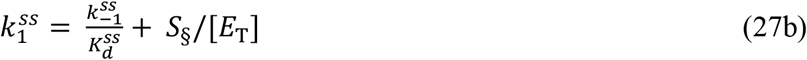

Equations (19) and (27b) are expected to give the same result. From either Eq. (23) or Eq. (25) is given:

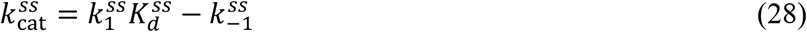

Then one can now substitute Eq. (27a) into Eq. (28) to give a composite equation such as:

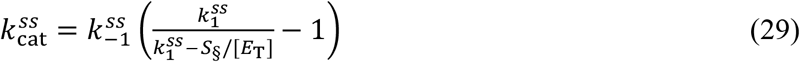

It should be constantly realised that steady-state shows linearity with time as the velocity of catalysis is directly proportional to substrate concentrations that are « *K*_M_; this experiment does not show the substrate concentration range that satisfies this requirement even if the substrate concentration range used for the assay is < *K*_M_ (the upper range is ≅*K*_M_/2). The variable *v* (the initial velocities, to be specific) should be proportional to such substrate concentrations.

### 2.3 Mutual dependence and contributing factors to the upper limits of rate constants, *k*_1_ and *k*_cat_

Meanwhile, there seems to be no direct way of calculating *k*_*-1*_, unlike *k*_*cat*_, which *is* given as *V*_max_/[*E*_T_]. What one can take home is that an increasing magnitude of *k*_cat_ results in a decreasing magnitude of *k*_-1_ and *vice versa*. It may appear that both parameters are mutually dependent, and factors that enhance *k*_cat_, such as the nature of the substrate, optimum ionic strength, optimum temperature, optimum pH, and the structure of the enzyme could compromise the magnitude of *k*_-1_ and *vice versa*. Going further demands that certain issues need to be pointed out clearly. This requires the scheme given below:

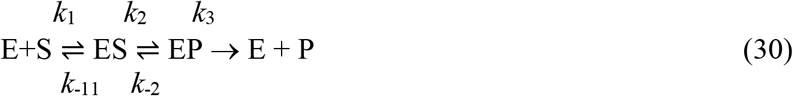

The overall events between the formation of ES, further conformational changes for catalytic function, bond breaking and formation in which molecular orbitals are involved, the chemistry or better yet the real biochemistry being the biological catalyst, leading to weak association between the product and enzyme, EP, and final release of the product, are defined as catalytic rate, *k*_cat_. More of the product is likely to be yielded if *k*_3_ + *k*_2_ **»** *k*_-11_ + *k*_-2_. The first-order rate constant, *k*_-11_ is introduced to emphasise that *k*_−1_ is the sum of the rate constants for the different stages leading to the dissociation of ES, the so-called “reverse reaction.” In the first place, there is no question of dissociation of nonexistent ES. Thus, all factors that increase and stabilise the transition state, E#S#, should ultimately increase the magnitude of *k*_cat_ and the much preferred catalytic efficiency, *k*_cat_/*K*_M_, because the ratio, the catalytic efficiency, *k*_cat_/*K*_M_ should be high at the expense of the “reverse” rate constant, *k*_-1_, for the process, ES→E+S. The stability of E^#^S^#^ is a function of the optimum pH, temperature, ionic strength, nature of the substrate, *etc*. While the life-span of E^#^S^#^ may be infinitesimal, an assumption of its nonexistence implies the absence of the chemistry of the process through which the enzyme ought to lower the “energy barrier,” thereby speeding up the reaction. Thus, the scheme (Eq. (19a)) is rewritten to show the importance role of E^#^S^#^.

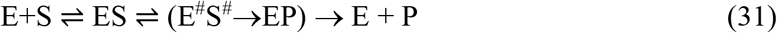

The process in brackets shows that there is no room for a backward reaction, due of course to the extremely high unfavourable energy barrier; it is either that the process, E^#^S^#^→EP,, occurs or the process, E^#^S^#^→ES, occurs. The process, ES⇌E^#^S^#^, is the preparation for catalytic function before product formation and the ultimate release of the product. It is only in the course of preparation that unfavourable perturbations, such as thermal, chemical, electronic perturbations *etc*, occur; such can lead to the process, E^#^S^#^→ES → E + S. According to equations (schemes) (19a) and (19b), the scheme E + S ⇌ ES → E + P described as unrealistic and unlikely is one in which the chemical transformation step and subsequent product release occur simultaneously with a single rate constant, *k*_2_ [14], which appears to be a summary of the entire catalytic events beginning with the noncatalytic event driven purely by diffusional encounter complex formation. The *k*_2_ in question is the *k*_cat_ in this research. One needs to realise that with increasing temperature, within the lower and optimum temperature ranges, the parameter *k*_cat_ increases, primarily as a result of the increasing rate of encounter-complex formation preceding ES formation.

Sometimes the interest of the pharmaceutical company and the medical community may be to control the biological function of one or more enzymes involved in the disease state of humans or animals. Drugs are manufactured and prescribed to treat such conditions, *e*.*g*., Parkinson’s disease; such drugs are likely to enhance the value of *k*_-1_ at the expense of *k*_cat_, and as is commonly observed, antidiabetic drugs are formulated and prescribed to inhibit the catalytic role of amylase.

Equating *k*_2_ to *k*_cat_ [1] can only be done under specified conditions and is not a likely phenomenon. Thus, *k*_*cat*_, being the overall rate constant, is governed by the magnitude of the net rate, depicting an imbalance between all forward reactions, with rate constants *k*_1_, *k*_2_, and *k*_*3*_, and backward reactions, with rate constants *k*_-1_ and *k*_-2_. No reaction proceeds in both forward and backward directions at the same time, as summarised in the above scheme, Eq. (19a). Rather, there are subpopulations of events, in which some ES break into free E and S while others elsewhere proceed to product. In between these events there is, characteristic of enzyme-catalysed reactions, a transition state depicted as ES ⇌ ES^#^ ⇌ EP; here is the key function of the enzyme, the lowering of the energy barrier, which is expectedly a very fast process. It is not certain whether, in a purely hydrolytic reaction, the breakdown of ES into E and S is enzyme catalysed, or, rather, whether it is a purely physico-biochemically oriented process.

It will require extraordinary mathematical formalism to enable a separate determination of the event, ES ⇌ E^#^S^#^ and E^#^S^#^ → EP; otherwise, if the durations of the processes, EP → E + P and E + S → ES, are known [15], then their rate constants can be determined. The duration of the process, ES → E^#^S^#^ → EP can also be calculated arithmetically. This analysis is important because other events and their rates affect the rate of the process, ES → E + S, and its rate constant. Establishing the value of *k*_-1_ in future research implies that the duration of all reverse reactions (EP → ES^#^ → ES) can also be determined. Thus, as was the case for the process, EP → P + E, whose duration and procedure for its determination were reported in the literature [15] (a part of which is awaiting an update), the duration of the process ES → E + S can also be determined. These may attract future investigation.

The issue about *k*_-1_ *vis-à-vis* the place of *V*_max_ or its equivalent, *k*_cat_, in the Michaelis-Menten constant equation needs clarification. To begin with, each aspect of the process, ES → E^#^S^#^ → EP, has its own duration, such that the sum of the reciprocals of all durations, including EP → E + P, yields the reciprocal of the catalytic rate, *k*_cat_. In addition, most enzymology authors use *k*_2_ as a first-order rate constant for EP → E + P; scheme (19a) addresses this issue in part. What plays out are various durations such as: *t*_ES_ (duration for the formation of ES), 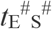 (duration of activated complex formation), *t*_EP_ (duration of EP formation after bond breaking and making), and *t*_*(EP*→ *E + P)*_(duration of product and enzyme release). Thus,

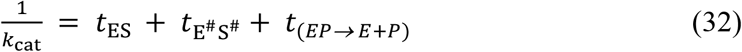

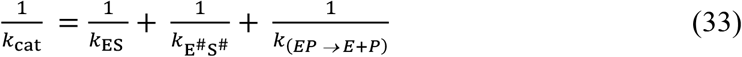

Similarly, each stage of the process has a time limit, E^#^S^#^ → ES → E + S. A direct process, EP → ES, may be a very remote possibility because the attachment of P to the active site may be purely physical, lacking capacity to assume a transition state complex. In this research, one should not be in doubt that alpha-amylase is not a synthase, which requires activated building blocks for the synthesis of starch through the Calvin-Benson cycle. The starting point is the conversion of glucose (Glc) to Glc-1-P with the concomitant conversion of adenosine triphosphate (ATP) to Adenosine diphosphate (ADP)–glucose (ADP-Glc), and pyrophosphate (PP_i_) by the catalytic action of ADP-glucose pyrophosphorylase (EC 2.7.7.27) [16]. Amylase has no place in biosynthesis, which requires the molecular fuel ATP to function. The equations for *t*_(EP→E+P)_ and *t*_ES_ have been derived elsewhere [15], but as previously stated, an update is required. Therefore, from Eq. (20), the following is derived:

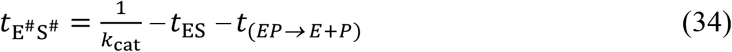

The implication is that:

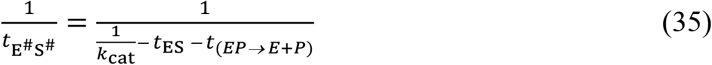

Thus, E^#^S^#^ → ES → E + S could be the scheme describing the reverse (backward) reaction with the following assay durations: 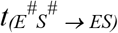 (deactivation of ES) and *t*_*(ES* → *E + S)*_ (breakdown of ES to E and S). The sum of the reciprocals of these durations gives *k*_-1_.

Hence,

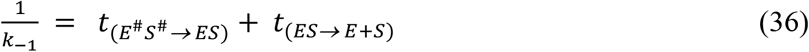

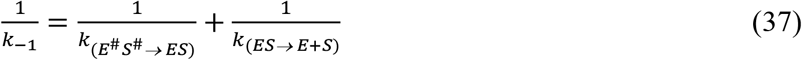

Most of the literature [3, 14] considers the rate constant *k*_-1_ to be the first-order rate constant for the process ES → E + S. It is assumed that E^#^S^#^ → ES → E + S is the overall rate constant for the entire process; if either *t*_*(ES* → *E + S)*_ or 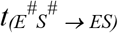 is derived and calculated, then either can be calculated. In this research, however, only qualitative treatment as described above is given for the duration of every aspect of the catalytic pathway. Quantitative treatment is reserved for the future. In summary, equations containing *k*_2_ must be replaced with equations containing *k*_cat_.

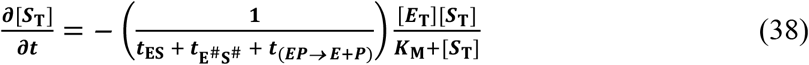

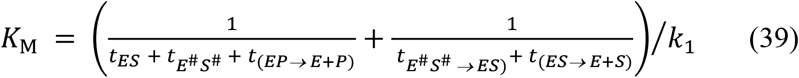

If one goes by the schemes, such as:

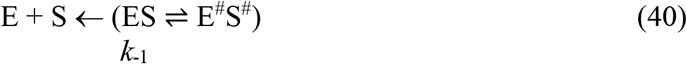

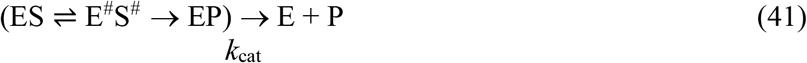

One may, then, consider two equilibrium dissociation constants, though one of them is rather known as Van Slyke-Cullen constant, *K*, such that:

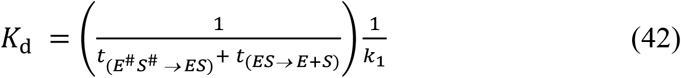

where *K*_d_ is the over-all dissociated constant at zero-order zone.

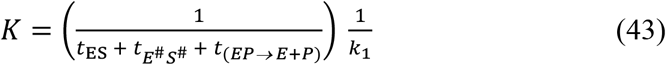

Therefore, in terms of time, the 2^nd^ order rate constant for the formation of ES is given as:

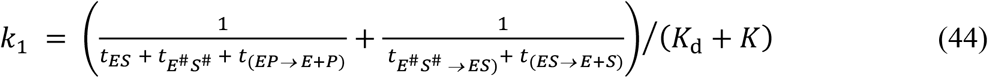

Recall too, that *K*_M_ = *K*_d_ + *K*.

## 3. MATERIALS AND METHODS

### 3.1 Materials

#### 3.1.1 Chemicals

*Aspergillus oryzea* alpha-amylase (EC 3.2.1.1) and insoluble potato starch were purchased from Sigma–Aldrich, USA. Tris 3, 5—di-nitro-salicylic acid, maltose, and sodium potassium tartrate tetrahydrate were purchased from Kem Light Laboratories in Mumbai, India. Hydrochloric acid, sodium hydroxide, and sodium chloride were purchased from BDH Chemical Ltd., Poole, England. Distilled water was purchased from the local market. The molar mass of the enzyme is ≈ 52 k Da [17, 18]. Distilled water was purchased from the local market. As a word of caution, readers of this paper should be aware that the use of the same enzyme in articles by the same author(s) is strictly due to budgetary constraints; however, this is not a serious concern because each paper addresses different issues, such as the evaluation of new models.

#### 3.3.2 Equipment

Electronic weighing machine was purchased from Wensar Weighing Scale Limited and 721/722 visible spectrophotometer was purchased from Spectrum Instruments, China; *p*H meter was purchased from Hanna Instruments, Italy.

### 3.2 Methods

#### 3.3.1 Preparation of reagents and assay

The method of assay of the enzyme is Benfield’s method [19] using gelatinised potato starch whose concentration range was 5-10 g/L. The reducing sugar produced upon hydrolysis of the substrate at 20°C using maltose as standard was determined at 540 nm with extinction coefficient equal to ≈ 181 L/mol.cm. The duration of assay was 3 min. A mass concentration = 2.5 mg/L of *Aspergillus oryzea* alpha-amylase was prepared in Tris HCl buffer at *p*H *=* 6.9; the choice of pH and temperature was at the authors discretion.

#### 3.1.2. The determination of rate constants

The pseudo-first order rate constant for substrate utilisation is determined as described in the literature [20], while the second order rate constant for the formation of the enzyme-substrate (ES) complex is determined as described elsewhere [15]. A double reciprocal transformation of the Michaelis-Menten equation [MM], otherwise known as the Lineweaver-Burk [21] plot, was used for the determination of the *K*_M_ and *V*_max_. All the steady-state parameters were determined as described in the theory section, where derivation and possible applications of derived equations can be found: Equations (18), (19), and Eq. (27a) are for the calculation of 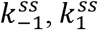, and 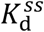 respectively.

### 3.3 Statistical analysis

Assays were conducted in triplicates. A method described by Hozo *et al*. [22] was used to determine the standard deviation (SD) for the median values, while Microsoft Excel was used to determine SD for the arithmetic mean values.

## 4. RESULTS AND DISCUSSIONS

There is no much data to analyse and discuss because the task or objective is to evaluate the derived equations and give evidence to the assumption that steady-state parameters are different in magnitude from zero-order kinetic parameters. There is also a need to come to the realisation that the chosen duration of the assay is pivotal to the calculation of kinetic parameters by any method: direct linear plot, double reciprocal plot (adopted in this research for convenience), nonlinear regression, *etc*. The substrate concentration comes next in importance; otherwise, if the duration of the assay is very long, an enzyme can be saturated even at a chosen lower substrate concentration range as long as such a range is > [*E*_T_]. This experiment yielded values of results beginning with what is considered the zero-order zone maximum velocity of catalysis, *V*_max_, the catalytic rate, *k*_cat_, and the Michaelian constant, *K*_M_ (Table 1). These kinetic parameters are needed for the calculation of steady-state constants according to Eq. (18). The same enzyme had been assayed in the past for different reasons, the determination of intrinsic rate constants [23, 24] under different conditions of temperature and pH, different concentrations of the enzyme, *etc*. This research, intended to quantify steady-state kinetic parameters, gave the 2^nd^ order rate constant for the formation of ES to be ≈ 4.781 exp. (+6) L/mol. min., or equivalently, 1.398 exp. (+4) L/g min. at a pH and ambient temperature of 6.9 and 20 °C, respectively, for [*E*_T_] ≈ 4.8077 M (Table 1) and [*S*_T_] ranging between 5 and 10 g/L. This differs from the previous report [24], which reported a value of 7.66 exp. (+6) L/mol. min. (or 2.24 exp. (+4) L/g. min.) for [*E*_T_] ≈ 3.205 exp. (−8) M; the *k*_cat_ values were expectedly different, with the current result being 1.599 exp. (+4) /min (Table 1), and the previous result being 2.108 exp. (+4) /min [24]. The *K*_M_ values were also different, being ≈ 18.65 g/L (Table 1) in this research and ≈ 37.24 g/L in the literature [24].

**Table 1:**
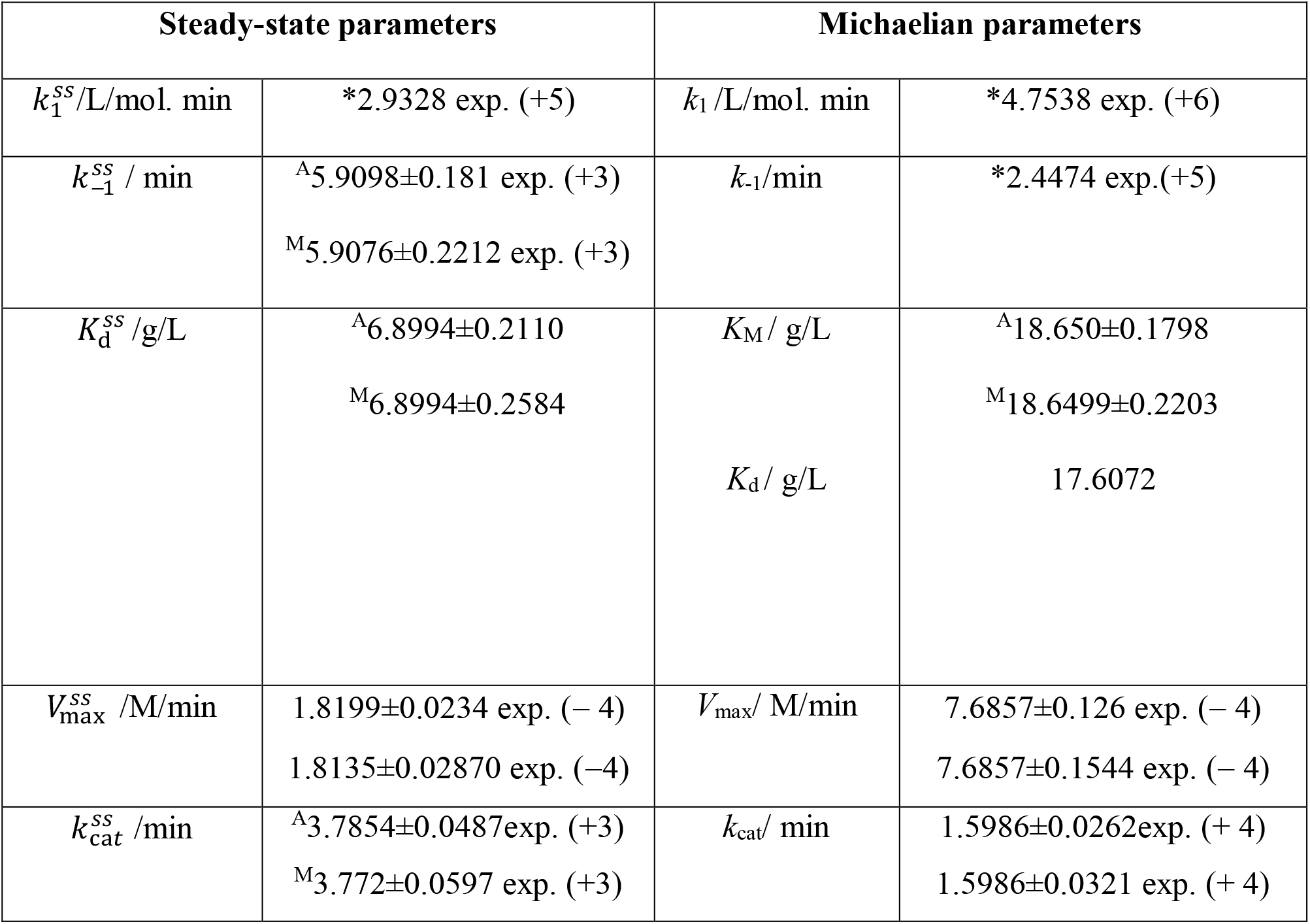
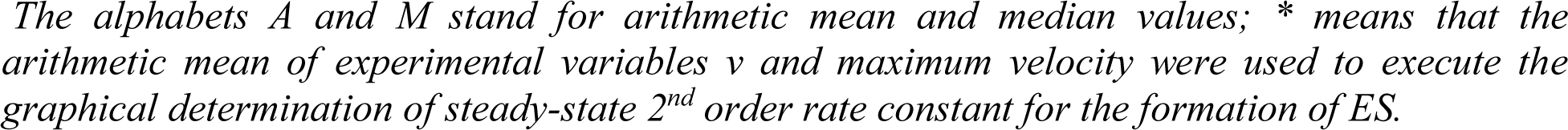
Steady-state and zero-order (Michaelian) kinetic parameters.

| Steady-state parameters |  | Michaelian parameters |  |
| --- | --- | --- | --- |
| $k_1^{ss}/\text{L/mol. min}$ | *2.9328 exp. (+5) | $k_1/\text{L/mol. min}$ | *4.7538 exp. (+6) |
| $k_{-1}^{ss} / \text{min}$ | <sup>A</sup>5.9098±0.181 exp. (+3)<br><sup>M</sup>5.9076±0.2212 exp. (+3) | $k_{-1}/\text{min}$ | *2.4474 exp.(+5) |
| $K_d^{ss} / \text{g/L}$ | <sup>A</sup>6.8994±0.2110<br><sup>M</sup>6.8994±0.2584 | $K_M / \text{g/L}$ | <sup>A</sup>18.650±0.1798<br><sup>M</sup>18.6499±0.2203 |
| | | $K_d / \text{g/L}$ | 17.6072 |
| $V_{\text{max}}^{ss} / \text{M/min}$ | 1.8199±0.0234 exp. (− 4)<br>1.8135±0.02870 exp. (−4) | $V_{\text{max}} / \text{M/min}$ | 7.6857±0.126 exp. (− 4)<br>7.6857±0.1544 exp. (− 4) |
| $k_{\text{cat}}^{ss} / \text{min}$ | <sup>A</sup>3.7854±0.0487exp. (+3)<br><sup>M</sup>3.772±0.0597 exp. (+3) | $k_{\text{cat}} / \text{min}$ | 1.5986±0.0262exp. (+ 4)<br>1.5986±0.0321 exp. (+ 4) |

*The alphabets A and M stand for arithmetic mean and median values; \* means that the arithmetic mean of experimental variables $v$ and maximum velocity were used to execute the graphical determination of steady-state 2<sup>nd</sup> order rate constant for the formation of ES.*

Because the literature is replete with concern for steady-state kinetic parameters such as the turnover number (*k*_cat_), the Michaelis-Menten constant (*K*_M_), and the specificity constant, *k*_cat_/*K*_M_ [1, 2, 5, 14, 25], the importance of sticking to what has been defined as zero-order kinetic parameters rather than the literature version cannot be overstated. This is notwithstanding the fact that the notion of steady-state assumption has been questioned in the past in a way that suggests its conceptual and operational inconsistencies. The ES that is part of equilibrium implies that it promotes a dynamic equation, but by so doing, it annihilates that equilibrium. The steady-state assumption implies a constant time-independent concentration (perhaps, d[ES]/dt ≈ 0) if [*S*_T_] is » [*E*_0_] in order to achieve dependence on other changing concentrations, but doing so invalidates constancy [26]. Despite the fact that this appears to be an ambiguous view in the literature, it is obvious that there could not be any result in the literature that suggests that zero-order kinetic values are different from steady-state values, except in this study. According to Table 1, the first-order rate constant for the dissociation of the ES to E and S is 5.908 exp. (+3) /min, which is significantly different from the zero-order value of 2.45 exp. (+5) /min. The difference is ≈ 97.59 % of the zero-order value. The steady-state catalytic rate constant (in contrast to a zero-order rate constant) as regarded in this research is ≈ 3.772 exp. (+3) /min, a value that is « than what used to be referred to as the asymptotic or zero-order value of ≈ 1.599 exp. (+4) /min; the difference is ≈ 76.41 % of the zero-order value. The steady-state 2^nd^ order rate constant, for the formation of ES is ≈ 2.933 exp. (+5) L/mol. min. Again, the zero-order value in this research is 4.781 exp. (+6) /L/mol. min. The difference is ≈ 93.87 % of the zero-order value.

The name, Michaelis-Menten constant reminds one that the zero-order parameters were instrumental to the derivation of the Michaelis-Menten equation. The constant is not intended to be equal to the steady-state dissociation constant 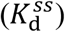, which is distinct from the Michaelis-Menten constant in this research; indeed, the value of 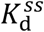 is ≈ 2.7-fold < the value of *K*_M_; however, this is a direct comparison; otherwise, the zero-order dissociation constant is given as: *k*_−1_/*k*_1_, where *k*_1_ is equal to 1.39 exp. (+4) L/g min, the equivalent of 4.7538 exp. (+6) L/mol. min, such that the result of computation gave result which is ≈ 17.61 g/L which is ≈ 2.56-fold > the value of 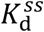 (Table 1).

Standing by the opinion held in a physical science research paper [27], this paper is not seen as sacrosanct and final; it may or it may not. This sentiment is anchored on the following premise: if the substrate concentration is « the *K*_M_, then the original Michaelis-Menten equation transforms to *v* = *V*_max_ [*S*_T_]/*K*_M_. If *v* is plotted versus [*S*_T_], and in particular [*E*_T_] is also > [*S*_T_], a linear relationship between *v* and [*S*_T_] is the case, with a possibility of the correlation coefficient being equal to one. Core kineticist and mathematical biologists in the subfields of enzymology and biochemistry are more familiar with asymptotes and the much-discussed hyperbolic curve relating velocities *v* versus [*S*_T_]. A straight line in any plot indicates that the dependent variable is directly proportional to the independent variable over a given time period. However, at much higher [*S*_T_], the linear relationship weakens; the magnitude of d[P]/dt, while increasing, is no longer proportional to the increasing availability of [*S*_T_].This is the result of the enzyme approaching and eventually reaching saturation when all of its molecules are participating in catalytic activities. Thus, maximum velocity is not achieved in the purely linear phase of the reaction characteristic of a steady state. It is always the case that *v* is plotted versus [*S*_T_], but different values of the former can be plotted versus different values of [*E*_T_] for each concentration of [*S*_T_] with a reasonable assay duration that must not lead to substrate depletion and saturation. In such a plot, a linear curve is expected with the coefficient of determination equal to one. What is very important is to realise that, if each substrate concentration is < the *K*_M_ for each value of [*E*_T_], one half of the maximum velocity of catalysis cannot be attained. Then how should an astute mathematical enzymologist known for very high standards and doing great work over the years explain why rate constants such as *k*_cat_ and the Michaelian constant, *K*_M_, are attributed to the steady-state phase, which is characterised by linearity, despite the fact that they are the product of nonlinearity in the hyperbolic relationship between initial rates and substrate concentrations?

## 5. CONCLUSION

Two assumptions stand out clearly as typical examples whose underlying principles are basically the same: They are the reactant stationary assumption and the quasi-steady-state assumption. All point to the need for the concentration of the enzyme to be less than [*S*_T_], let alone (*K*_M_ + [*S*_T_]); yet the view that the steady-state assumption can be valid without ensuring [*S*_t_] ≈ [S_0_] during the initial transient for the reaction mechanism depicted by the scheme, E + S ⇌ ES → E + P, appeared to question the concept of zero-order kinetics. If Δ[S]/[*S*_T_] is not ≈ zero, it is likely that d[ES]/d*t* will not be ≈ zero because it is the abundance of S in excess of E, that ensures the recombination of E with S. The fact that *v* = [ES]/*k*_cat_ implies that [ES] remains incalculable if information about zero-order kinetics (a nonlinear phenomenon) and its kinetic parameters are unknown. The initial variable is a fractional part of an asymptotic rate of catalysis as expressed by *V*_max_ [*S*_T_]/([*S*_T_ + [*S*_T_]), because [*S*_T_]/([*S*_T_ + [*S*_T_]) is < 1. The summary is that *V*_max_ is not attained where [*S*_T_] is < *K*_M_. The steady-state equation for the calculation of the first-order rate constant for the dissociation of ES into E and S was derived and calculated. The calculated steady-state first-order rate constant was « the zero-order, Michaelian value, and the difference is ≈ 97.59 % of the zero-order value; the steady-state catalytic rate differed from the zero-order catalytic rate by ≈ 76.41 % of the latter value; and it was ≈ 93.87 % with respect to the second-order rate constant for the formation of ES. Rates as a function of time were derived, leaving the possibility that future research will focus on a method for the determination of the duration and consequently the rate of the process, ES → E + S, distinct from the process, E^#^S^#^ → ES.

## Supporting information

RE-SS-SUB...Steady-state reverse rate constant and reaction pathway rate constants..(Autosaved) (6).pdf-Adobe reader)

## ACKNOWLEDGMENTS

The supply of electricity by the Management of Royal Court Yard Hotel, Agbor, Delta State, Nigeria during the preparation of the manuscript is deeply appreciated.

## AUTHOR’S CONTRIBUTIONS

The sole author designed, analysed and interpreted and prepared the manuscript.

