## Supplementary figures and images for "Derivation of Steady-State First-order Rate Constant Equations for Enzyme-Substrate Complex Dissociation, as well as Zero-order Rate Constant Equations in Relation to Background Assumptions"

### RE-SS-SUB...Steady-state reverse rate constant and reaction pathway rate constants..(Autosaved) (6).pdf-Adobe reader)

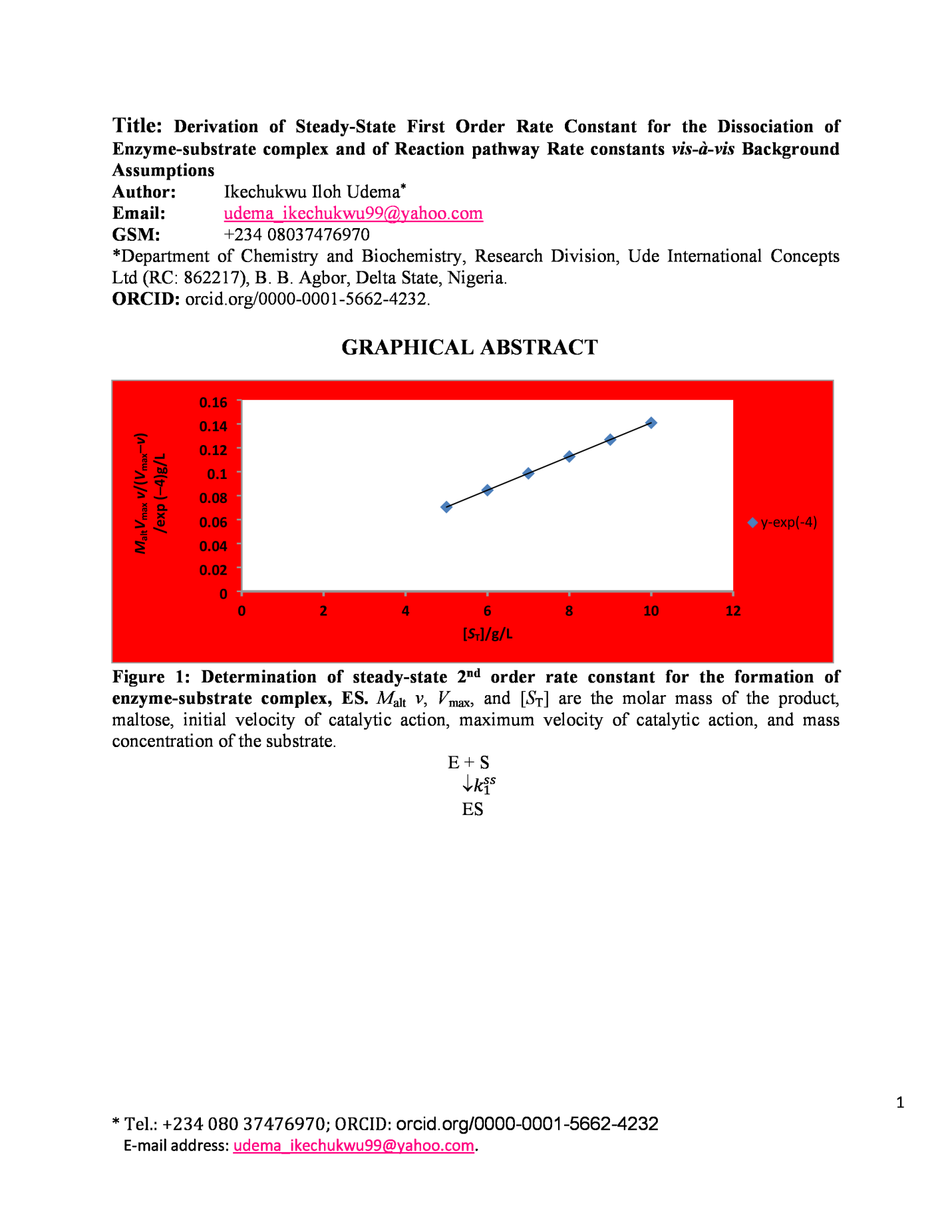

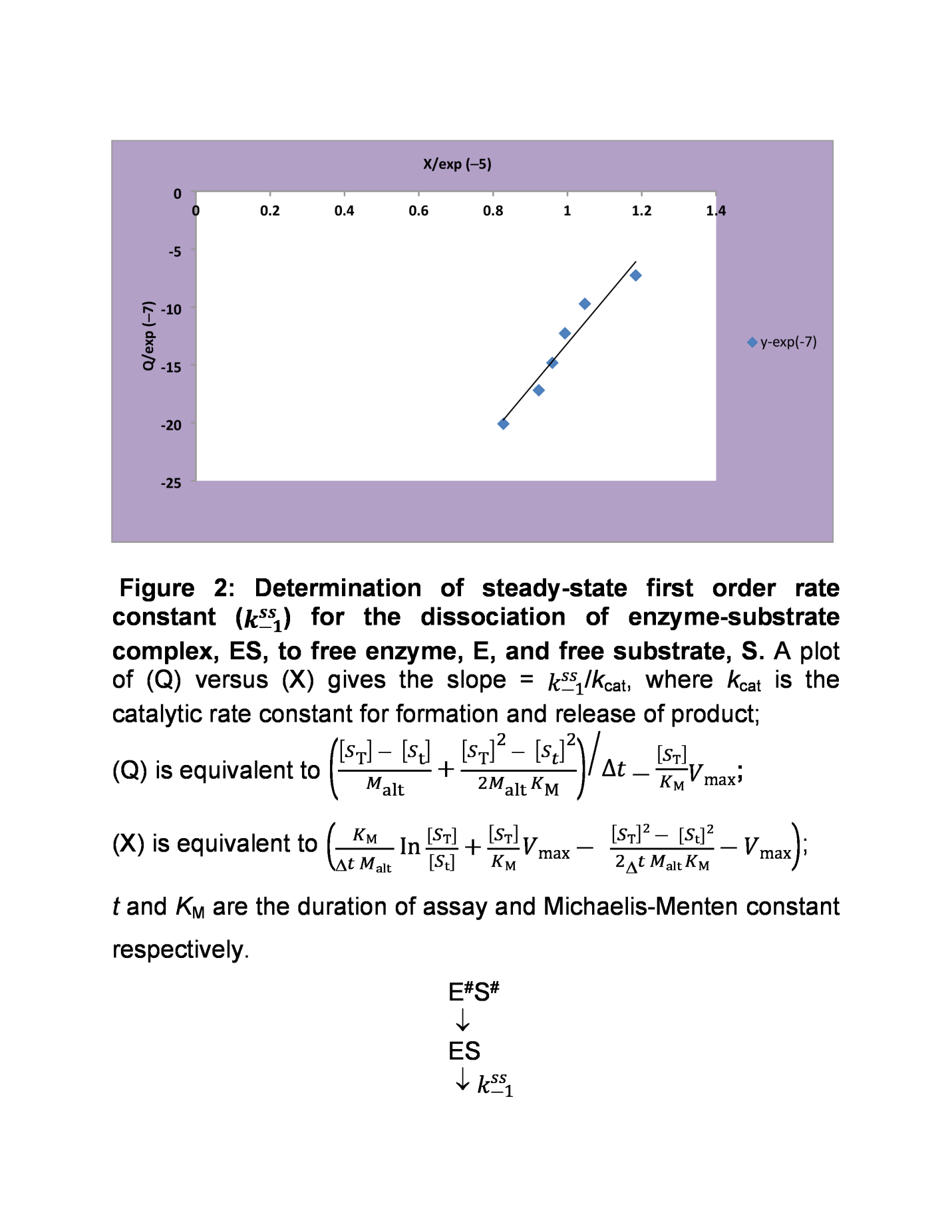

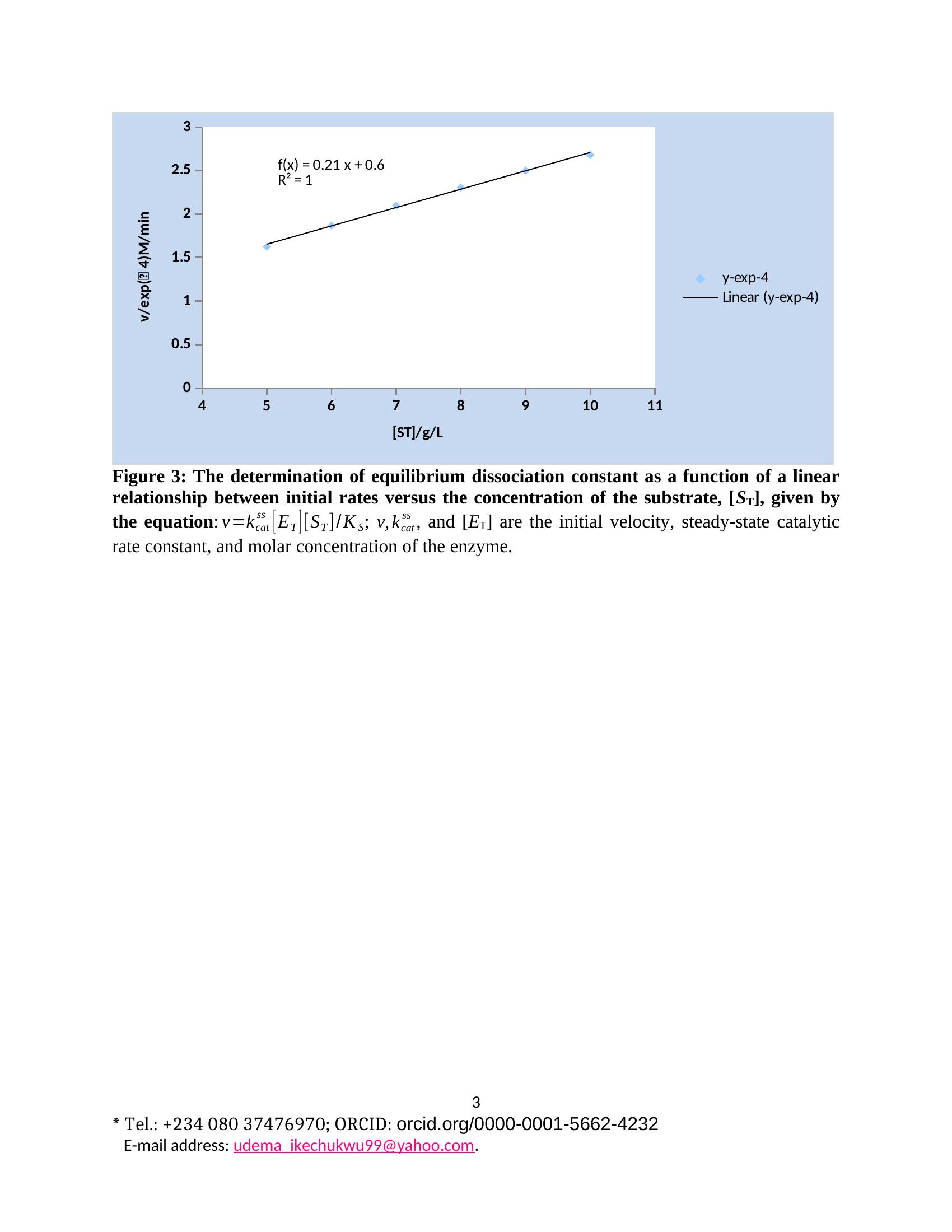
